# Apical Actin-myosin Network Regulates the Tight Junction of Polarized Madin-Darby Canine Kidney Cells

**DOI:** 10.1101/2021.05.30.446323

**Authors:** Chia-hsuan Lu, Fu-Lai Wen, Shawn Ching-Chung Hsueh, Wen-hsiu Wu, Yu-Fang Lin, Mathieu Prouveur, Thomas Boudier, Keng-hui Lin

**Affiliations:** Department of Computer Science and Information Engineering, National Taiwan University, Taipei, Taiwan; Institute of Physics, Academia Sinica, Taipei, Taiwan; Department of Physics, University of British Columbia, Vancouver, Canada; Department of Physics, National Tsing-hua University, Hsinchu, Taiwan; Department of Electrical Engineering, National Taiwan University, Taipei, Taiwan; Department of Applied Mathematics, Mines ParisTech, Paris, France; Institute of Molecular Biology, Academia Sinica, Taipei, Taiwan; Sorbonne University, Paris, France; Department of Physics, National Taiwan University, Taipei, Taiwan

## Abstract

The tight junction outlines the apicolateral border of epithelial cells like a belt, sealing the paracellular space when cells form contacts with each other. The permeability and morphology of tight junction are regulated by actomyosin contractility, which has been conventionally thought from the purse-string-like circumferential actomyosin belt along tight junction. Spatially, the tight junction is close to the apical actin network, which exerts inward contractions orthogonal to the tight junction. To test the contributions from apical actin network, we laser-ablated spots on the apical surface of polarized Madin-Darby Canine Kidney (MDCK) epithelial cells. Laser ablation severed the apical cytoskeleton network, decreased in-plane tension, increased the apical surface area, and rendered the tight junction less tortuous in shape. Consistent with these observations, changes in MDCK cell sheet morphology due to cell proliferation, or perturbation with the ROCK inhibitor Y27632 increased the density of the apical actin network and decreased tight junction tortuosity. The morphological analysis revealed scutoids in flat MDCK cell sheets, contrary to predictions from a previous model that only considered cell-cell interactions as line tension. Additional cell-cell interactions from apical in-plane tension provides probable cause for the occurrence of scutoids on flat geometry. Taken together, our findings identify the importance of the apical actin network exerting in-plane apical tension to regulate tight-junction mechanobiology and epithelial cell shape.

**Significance Statement:** The tight junction is located at the apicolateral cell border and regulates paracellular diffusion. Adjacent to the tight junction, the actin cytoskeleton forms a dense network beneath the apical surface and an actomyosin belt that circumscribes the lateral surface of the cell. Tight junctions are connected to the actin cytoskeleton which regulates paracellular transport, but the role of tension-mediated regulation of the tight junction by various actin structures is poorly understood. Here, we provide evidence that tension on the tight junction is mediated by the apical actin network. Our results provide a reinterpretion of past reports and broaden our understanding the mechanobiology of tight junctions.

## Introduction

Epithelial cell sheets shape the embryo and line the surfaces of many organs in adults. They act as barriers that compartmentalize biological spaces; the three sides of the plasma membranes of epithelial cells (apical, basal, and lateral surfaces) face different compartments (lumen, extracellular matrix, and neighboring cells, respectively; Fig. 1A). Changes in the interfacial tensions of these three surfaces govern tissue morphogenesis, motivating developmental biologists and cellular biophysicists to unravel the pertinent mechanisms. Recently, theoretical biophysicists have made predictions about cell shape and sheet structure based on the relative interfacial tensions of these three surfaces (1, 2), explaining some observed tissue patterns (3) and predicting a new shape, the scutoid, in epithelia (4). Further understanding of morphogenesis requires detailed characterization of the tension-generating mechanisms at play in cellular surfaces.

**Fig. 1.**
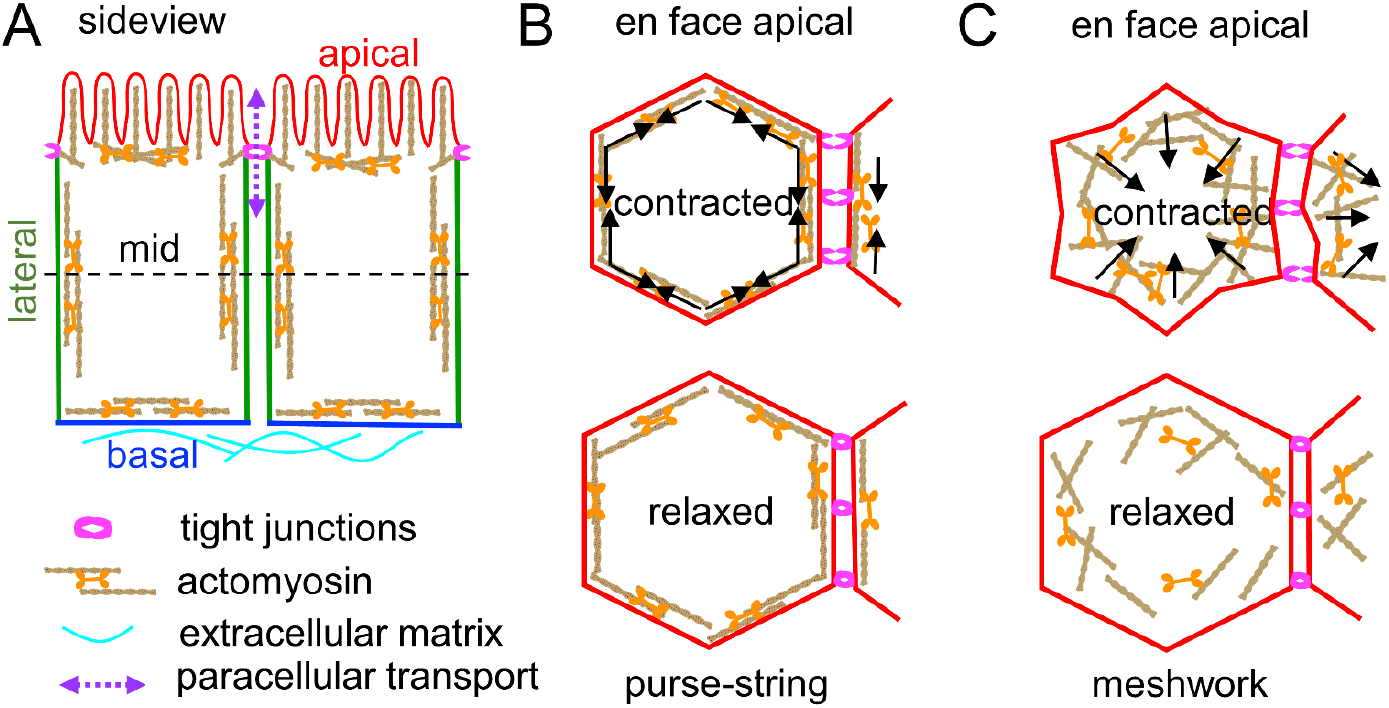
Schematics of actomyosin organization in epithelial cells and how contractility affects the paracelluar transport of tight junctions. (A) Sideview of polarized epithelial cells. Tight junctions at the apicolateral border seal the paracellular space, causing transport to occur through the tight-junction pores. (B) En face apical view of the purse-string model. Contraction along the circumferential actomyosin belt constricts apical areas and opens the pores of tight junctions at the borders of the apical surface. (C) En face apical view of in-plane apical tension. Medial contraction constricts apical areas, wrinkles tight junctions, and opens the pores of tight junctions at the borders of the apical surface.

One of the major events during morphogenesis is epithelial folding and tube formation via apical constriction: the apical side of an epithelial cell shrinks, leading to bending of a cell sheet and formation of a hollow tube. The most prevalent explanation for apical constriction is the purse-string model (5, 6), which posits that F-actin is bundled into a circumferential belt on the apical plane and that actomyosin contraction reduces the apical area relative to the rest of the cell (Fig. 1B). However, a growing body of evidence indicates that apical constriction can be driven by other force-generating mechanisms, such as pulsed medial actomyosin contraction in *Drosophila* morphogenesis (7, 8) (Fig. 1C).

The tight junction is localized at the border between the apical and lateral surfaces of epithelial cells (Fig. 1A) and is associated with the actin cytoskeleton (9). The tight junction delimits the boundary of the apical surface and controls paracellular diffusion. Tight junction morphology and function were previously proposed to be regulated by the purse-string contraction of the actomyosin belt (10). The actin-binding protein Shroom, which is required for neural tube closure (11-13), regulates the apical tension of Shroom-expressing Madin-Darby Canine Kidney (MDCK) cell sheets (a widely studied *in vitro* epithelial cell model) (14, 15) to have a smaller apical surface area and straighter tight-junction morphologies than wild-type cells (11). In Shroom-expressing cells, myosin IIB colocalizes with the tight junction protein ZO-1 and displays sarcomeric structures along the tight junction (11). These observations based on Shroom-expressing cells corroborate the purse-string model for regulating tight-junction organization (Fig. 1B).

The purse-string model was also proposed to explain the increase in permeability of tight junctions via the opening of tight-junction pores by increasing the contractility of the circumferential actomyosin ring as line tension (Fig. 1B) (10). Indeed, increases in actomyosin contractility and the tortuosity and permeability of tight junctions have been reported in MDCK cells devoid of Shroom (16, 17) and Caco-2 human cells (18). However, high line tension favors shorter and linear tight-junction morphologies, in contrast to the tortuous tight junction morphologies observed previously (16-18). Further, although the tight junction is known to be regulated by actomyosin contractility, it has not been associated with force transmission (19). In contrast, E-cadherin junctions, a major cell-cell junction located at the lateral epithelial surface that connects to the circumferential actomyosin belt, are well-described by the purse-string model as the main route of force propagation to neighboring cells (20). E-cadherin junctions and tight junctions together form the apical junction complex (21); thus, the dominance of E-cadherin junctional tension could cause the contributions of in-plane apical tension in tight junction structure and function to be overlooked.

To close these conceptual gaps, we hypothesized a different force-generating direction for apical tensions exerted on the tight junction in polarized MDCK cell sheets. We investigated apical actomyosin networks, rather than the circumferential actomyosin belt, which generate in-plane surface tensions across the apical surface onto the tight junction (Fig. 1C). We used laser-ablated on the apical surface of polarized MDCK cells and a semi-automatic image-processing workflow that includes cell segmentation, edge extraction with subpixel fit for accurate contour length computation, and vertex extraction to quantify MDCK cell shapes. When fewer apical actin cytoskeletons were present under perturbations including laser ablation, changes in cell density, and actomyosin inhibition, tight junctions were less tortuous and apical surface areas were larger than in unperturbed cells. Overall, our results indicate that apical tensions arise from the apical actin cortex network through the tight junction, raising questions about previous models that only considered tensions on the tight junction exerted by the circumferential actomyosin belt.

## Results and Discussion

### Laser ablation at a spot on the apical surface leads to area expansion

Laser ablation, which has been widely used to quantify mechanical tension in living organisms (8, 22, 23), disrupts the local contractile apparatus, thereby releasing contractile forces and causing morphological changes. In order to visualize such apical changes in MDCK cells, we engineered cells to express a fluorescently labeled tight-junction protein (EGFP-ZO-1; see Materials and Methods) and grew the fully polarized cells on filters for 3 days, at which point tight junctions became tortuous (Fig. 2A, green). The EGFP-ZO-1 signal delimited the edge of the apical surface and was used to measure changes in the contour and hence the area of the apical surface after laser ablation. A pixel-width square on the apical surfaces of these MDCK cells was ablated (Fig. 2A, cross). Most cells underwent an expansion in apical surface area (Fig. 2A, magenta). Interestingly, some edges moved more than other edges, indicating that this expansion was not uniform in all directions. In addition, we detected a gradual decrease in the tortuosity of the tight junction after laser ablation (Fig. 1E). These morphological changes indicate that a heterogeneous contractile network at the apical plane causes the tortuous shape of tight junctions and controls the apical surface area of the cell sheet. Laser ablation also prompted the movement of cells beyond their nearest neighbors (Movie S1), indicating long-range connections of the apical network between cells, perhaps through tight junctions.

**Fig. 2.**
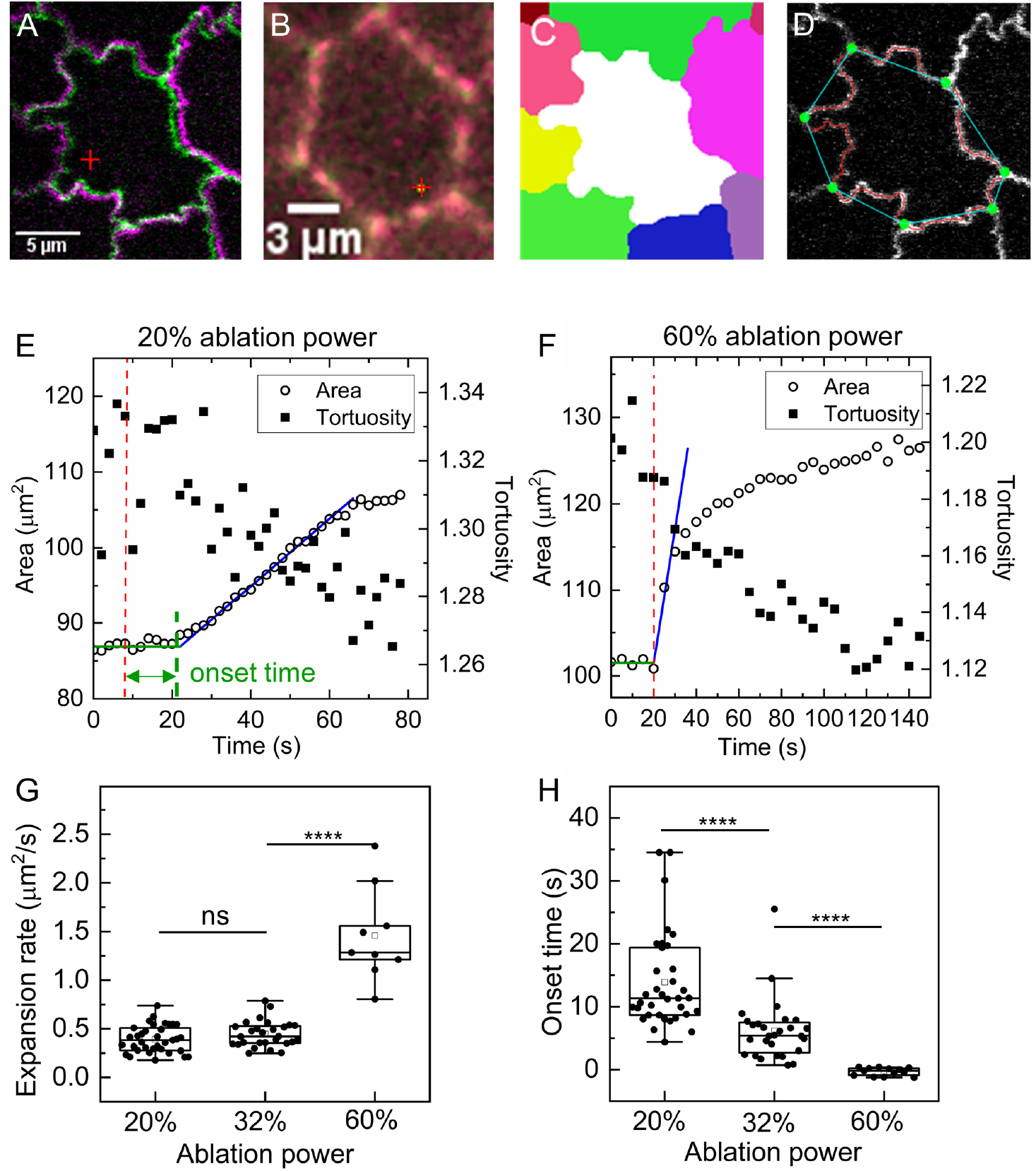
Laser ablation on the apical surface of MDCK cells induces an increase in apical area and decreases in tight-junction tortuosity. (A) Overlaid confocal images of the apical surfaces before (green) and 1 min after (magenta) ablation. The apical area of the post-ablation cell visibly increases. The ablation spot is marked with a cross. (B) Overlaid images of the mid-section before (green) and 1 min after (magenta) ablation. There is no change in cell shape. The ablation spot is marked with a cross. (C) Segmented cells based on a watershed approach (Materials and Methods). (D) Sub-pixel-resolved contours (red) enable calculation of contour perimeters and the vertices (green dots) of a cell to compute the polygonal perimeter (cyan). Tortuosity is the ratio of the contour perimeter to the polygonal perimeter. (E) Apical area increases and tortuosity decreases over time after ablation at 20% laser power (red dashed line). The expansion rate is calculated based on fitting the fast-expanding part of the cell (blue line; Materials and Methods). Onset time is the time interval from ablation (red dashed line) to the cross-over point of the baseline (green line) and the expansion fitting line (blue line). (F) Apical area increases sharply and tortuosity decreases when cells are ablated (red dashed line) at 60% laser power. (G) Average expansion rates at 60% laser power are significantly higher than average expansion rates at 32% and 20% power (*p* < 10^−6^). (H) Onset time increases significantly when ablation power decreases. The symbols, ns and 4 asterisks, denote *p* ≥ 0.05 and *p* ≤ 10^−4^, respectively.

The mid-section (a single z-slice around 50% of cell height, Fig. 1A) of MDCK cells served as a control to isolate the effects of laser ablation on the circumferential actomyosin belt because it localizes to the lateral surface alone, as described by the purse-string model (20). When we laser-ablated a pixel-width box in the mid-section of MDCK cells expressing a fluorescently labeled F-actin binding protein (GFP-F-tractin; Materials and Methods), the cell shape remained unchanged (Fig. 2B). Even when we enlarged the ablated region into a line, the cell shape did not change after ablation (Movie S2). Since there is no actomyosin network in the middle of the mid-plane to be ablated, cell shape does not change upon ablation *per se*. If the tight junction is mainly connected to the circumferential belt, then there should be no change in cell shape after ablation, as observed only in the mid-section of the cell.

### Expansion dynamics depend on ablation power

In order to characterize the mechanisms underlying apical-surface expansion after ablation at the apical cell surface, we carried out quantitative image analysis of morphological changes of these cells after ablation at various laser powers. Briefly, we computationally segmented each apical surface contoured by tight junctions and measured its apical area (Fig. 2C; Material and Methods) and tortuosity, which is defined as the ratio of the contour perimeter (Fig. 2D, red curve) to the polygonal perimeter (Fig. 2D, cyan polygon; vertices shown as green dots; SI Materials and Methods). The original area and tortuosity of the cell are averaged from the first five frames before ablation. We only analyzed the expansion rate of cells expanding >10% over the original area. Most cells survived laser ablation at a pixel-width square when laser power was ≤ 32% (Fig. S1A). Higher laser power and larger regions of ablation led to increased cell death and cell extrusion (Fig. S1B).

The percentages of cells that expanded after ablation were 100%, 85%, and 66% for ablation at 60% (*N* = 14), 32%, (*N* = 39), and 20% (*N* = 64) power, respectively. After ablation, apical surface areas increased and tight junction tortuosity decreased (Fig. 2E). We defined the expansion rate as the maximum slope from a cell that underwent sustained expansion (Fig. 2E, blue line; Materials and Methods). Cells ablated at 32% power displayed a similar expansion curve as cells ablated at 20% power (Fig. 2E). Usually there was a delay before a nearly constant and sustained expansion in apical surface area ensued; onset time was defined by the time from ablation to the crossover between the initial baseline (Fig. 2E, green) and the expansion line (Fig. 2E, blue). In contrast, cells ablated at 60% power exhibited a sudden increase in surface area immediately after ablation (Fig. 2F). The average expansion rate, 1.46 ± 0.48 μm^2^/s, after ablation at 60% power (*N* = 14) was much larger than the average expansion rates, 0.44 ± 0.13 μm^2^/s (*N* = 28) and 0.39 ± 0.14 μm^2^/s (*N* = 35), at 32% and 20% power, respectively (Fig. 2G). While there was almost no onset time at 60% power (Fig. 2H), onset time increased significantly from 6.2 ± 4.9 s at 32% laser power (*N* = 28) to 13.9 ± 7.6 s at 20% power (*N* = 35) (Fig. 2G). The slow onset of constant expansion of apical surface area after ablation indicates that expansion at 20% and 32% power is not driven by fast elastic recoil, whereas the instant expansion at 60% is likely due to recoil (23).

In order to shed light on the mechanism underlying post-ablation cell expansion, we sought additional factors upon which expansion rates depended. Interestingly, at 20% and 32% power there was a strong and significant correlation between expansion rate and original apical surface area before ablation (Fig. S2A), but a weaker correlation between expansion rate and original tight-junction tortuosity before ablation (Fig. S2B). In comparison to low-power ablation, the expansion rates of cells ablated at 60% power correlated not only with original apical surface area (Fig. S2C) but also with tight junction tortuosity (Fig. S2D). The immediate recoil-based expansions after high-power ablation should depend on pre-existing apical tension, which underlie the original tight-junction tortuosity of the cell. We speculate that the correlation between expansion rates and original apical surface areas arises from the removal of apical forces from the ablated cell, and that these apical forces are proportional to the original apical surface area of the cell.

### Ablation reduces F-tractin levels at the apical surface

Since we observed non-recoil expansion of apical area after low-and median-power ablation, we sought to understand the effect of ablation on the apical actin cytoskeleton network, which generates and transmits tension. In order to visualize F-actin, we ablated MDCK cells expressing F-tractin-EGFP and inferred F-actin dynamics based on changes in F-tractin localization (Materials and Methods). Interestingly, the confocal *en face* apical maximum-projected image sequences revealed that the overall F-tractin signal decreased over time after laser ablation at 32% power (Fig. 3A, Movie S3). Some of the punctate F-tractin signal in microvilli became less pronounced than before ablation, but they persisted after laser ablation at 32% power (Fig. 3A). A sideview scan revealed that after laser ablation at 32% power, apical F-tractin appeared diffusely in the cytosol (Fig. 3B, Fig S3).

**Fig. 3.**
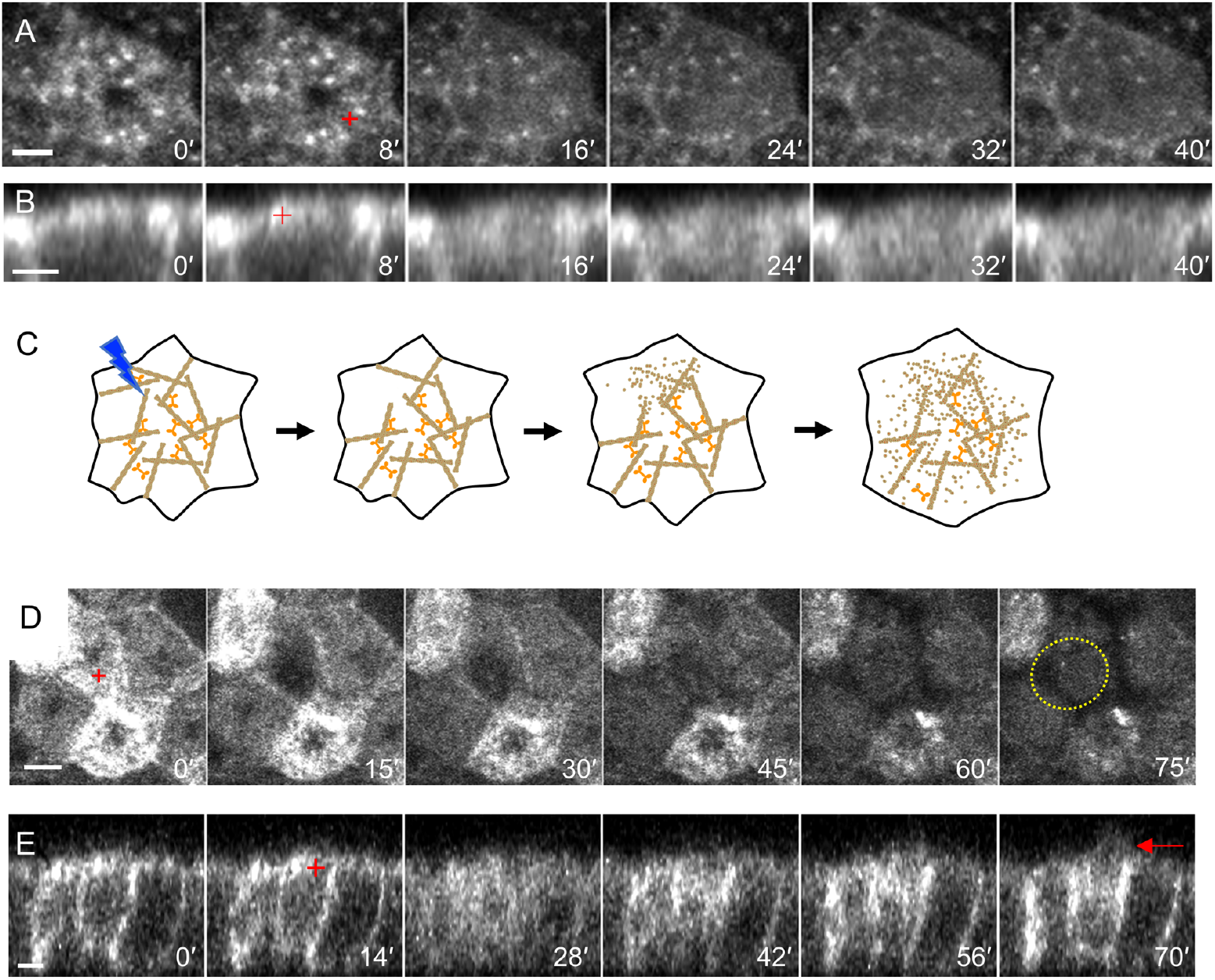
Dynamics of F-tractin after ablation indicate a decrease in the number of F-tractin molecules at the apical surface. The red cross marks the location of ablation at 32% power (second frame in panels A and B). Scale bars, 3 μm. (A) Confocal en face apical maximum-projected image sequences show that F-tractin signals decrease over time after ablation. (B) *xz* scanning confocal image sequences indicate that apical F-tractin molecules descend into the cell body after ablation. (C) Proposed mechanism explains the decrease in F-tractin signal due to cascading severing of the F-actin network by actin depolymerization factors, removing local tension. (D) Confocal en face apical image sequences indicate that F-tractin signals decrease dramatically after ablation at 60% power. The red cross marks the location of the ablation. Scale bar, 5 μm. The yellow dotted circle highlights the formation of a blob in the last frame. (F) *xz* scanning confocal image sequences reveal that apical F-tractin molecules are extruded from the cell (red arrow). Scale bar, 3 μm.

We postulate that at lower laser powers (32% and 20%), apical F-tractin is severed by ablation, resulting in apical F-tractin diffusing into the cytoplasm. Hayakawa *et. al*. previously reported that a suspended actin filament *in vitro* was severed in 17 ± 3.9 s after cofilin application, and when the actin filament was under tension, the severing by cofilin was significantly delayed (43.2 ± 3.9 s) (24). Therefore, we speculate that laser ablation removes a small local region of the apical actin network, thereby removing local tensions and initiating a cascade of actin severing (Fig. 3C). The effect of laser power on apical surface area expansion rate and on the onset time of that expansion could be also explained by this hypothesis. The larger the region of actin network destroyed by the slightly higher power laser, the more actin that is severed; apical surface area expansion is not pronounced until a substantial amount of actin is severed, manifesting as an onset time for expansion. Since expansion is due to actin severing, expansion rates do not depend on the tortuosity of the tight junction (Fig. S2B, right).

Confocal *en face* image sequences showed that the F-tractin signal of laser-ablated cells decreased significantly after ablation at high (60%) laser power (Fig. 3D); the F-tractin signal in neighboring cells also decreased (Fig. 3D). Approximately 1 min after laser ablation, a faint amorphous blob of F-tractin appeared around the ablation site (Fig. 3D, yellow circle). Side-view *xz* scans also indicated that apical F-tractin signal decreased in both ablated and neighboring cells (Fig. 3E). Here we clearly detected F-tractin extruded from the cell surface ∼1 min after ablation (Fig. 3E), consistent with the *en face* images (Fig. 3D). This extrusion may explain the two-stage change in the area-expansion curve after ablation (Fig. 2F): the initial fast expansion is due to elastic recoil and the second slow expansion is due to extrusion of apical actin network (Fig. 2F).

Overall, our data show that at both high and low laser powers, F-tractin levels at the apical surface decrease through two distinct processes: a direct mechanical disruption at high power and a disassembly reaction induced at low power. Although we did not directly observe cellular F-actin in these studies, a reduction in the presence of the F-actin binding protein F-tractin can also result in a reduction in actin cortical tension (25), leading to increases in apical area and decreases in tight junction tortuosity.

### Tight junction tortuosity is correlated with apical F-actin levels

These results prompted us to investigate the correlation between apical area, tight junction tortuosity, and apical F-actin levels. To this end, we performed live-cell imaging of MDCK cell sheets from day 2 to day 3 (Fig. 4A, Movie S4) after initial seeding of a culture dish at 600 cells/mm^2^ on day 0 (Materials and Methods). The mean apical surface area decreased from 170.6 ± 53.6 μm^2^ (day 2, *N* = 115) to 134.2 ± 39.4 μm^2^ (day 3, *N* = 163) in the same field of view (Fig. 4B). Notably, the morphology of tight junctions changed dramatically from relatively straight on day 2 (τ = 1.06 ± 0.02) to tortuous on day 3 (τ = 1.22 ± 0.09) (Fig. 4C).

**Fig 4.**
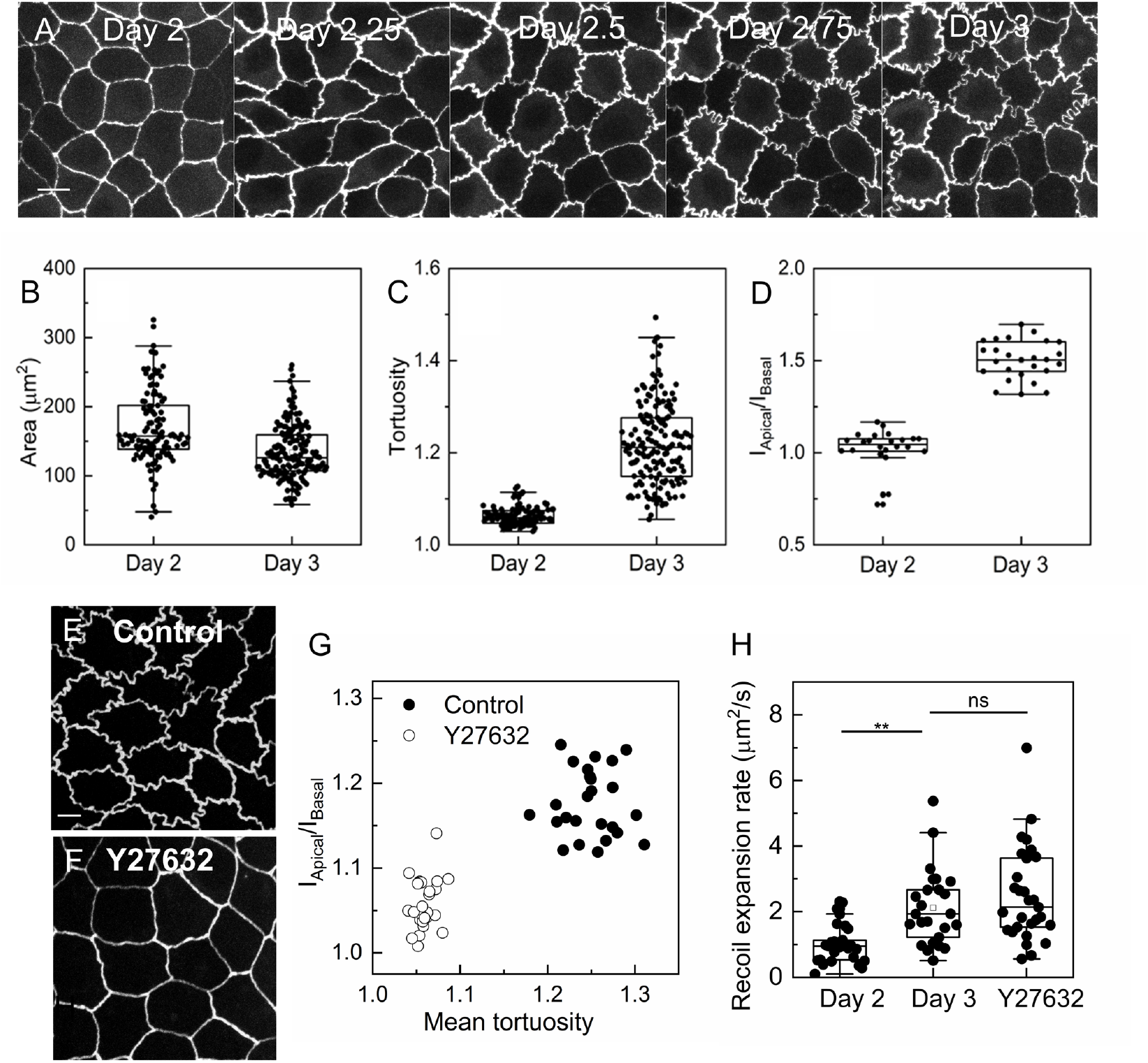
Changes in cell area, tortuosity, the apical/basal F-actin intensity ratio, and recoil expansion rates of cells grown for 2 days, 3 days and under Y27632 treatment. (A) Time-lapse montage of the tight junction of an MDCK cell sheet from day 2 to day 3 after seeding on day 0. (B, C) Apical areas decrease significantly (B; *p* = 6×10^−9^) and tortuosity increases significantly (C; *p* = 4×10^−45^) in MDCK cell sheets on days 2 (*N* = 115) and 3 (*N* = 163) in the same field of view depicted in (A). (D) The increase in apical F-actin from day 2 (*N* = 26) to day 3 (*N* = 24) is deduced from the F-actin intensity ratio of the apical surface to the basal surface (*p* = 2×10^−14^). (E, F) Micrographs of the tight junction-associated reporter EGFP-ZO1 expressed in an MDCK cell sheet under (E) control conditions and (F) after treatment with the ROCK inhibitor Y27632. Scale bar, 5 μm. (G) Ratio of F-actin intensity in the apical plane to that in the basal plane vs. the mean tortuosity of all cells in one image stack. (H) Recoil expansion rates of MDCK cells grown for 2 days, 3 days, and after treatment with Y27632. The symbols, ns and 2 asterisks, denote *p* ≥ 0.05 and *p* ≤ 10^−2^, respectively.

Hallmarks of MDCK cell polarization include the establishment of tight junctions and increases in the number of microvilli emanating from the apical membrane; each microvillus contains a dense bundle of F-actin, which can be used as a surrogate marker for the microvillus at the apical surface (26, 27). When we quantified the ratio of F-actin intensity at the apical and basal planes (*I*_apical_/*I*_basal_; Fig. S4; SI Materials and Methods), we detected an increase in this ratio from day 2 to day 3 (Fig. 4D). We speculate that the more-concentrated F-actin meshwork associated with the increased numbers of microvilli generate higher actomyosin contractility and transmitted more force than the less-dense F-actin network associated with less well-polarized cells containing fewer apical microvilli (25, 28). Such a relationship between apical actin concentration and tight junction tortuosity based on cell proliferation is consistent with the observation based on laser-ablation experiments.

Previous studies demonstrated that tight junctions straighten when MDCK cell sheets are treated with the ROCK inhibitor Y27632 (16). We therefore quantified mean tortuosity and *I*_apical_/*I*_basal_ of F-actin signals on images of untreated MDCK cell sheets and Y27632-treated MDCK cell sheets (Fig. 4E, F). Tortuosity positively correlated with *I*_apical_/*I*_basal_ (Fig. 4G).

To compare the level of medio-apical tension in the apical surface under different treatment, we measured the recoil expansion rate by ablation at high power. We ablated the apical surface with a diffraction-limited spot using an LSM 980 laser at 50% ablation power. The expansion dynamics of the cells ablated by LSM 980 (Fig. S5) were similar to the expansion dynamics of cells ablated at 60% power with an LSM 510 laser (Fig. 2F): a rapid increase in area immediately after ablation followed by a gradual increase in surface area (Fig. S4). The recoil expansion rate was calculated by taking the slope between the time points immediately before and after laser ablation. The recoil expansion rates for untreated MDCK cells on day 2, untreated MDCK cells on day 3, and MDCK cells treated with Y27632 were 1.02 ± 0.56 μm^2^/s (*N* = 29), 2.12 ± 1.16 μm^2^/s (*N* = 25), and 2.5 ± 1.44 μm^2^/s (*N* = 29), respectively (Fig. 4H). The increase in recoil expansion rate from day 2 to day 3 was consistent with our expectations; apical tension increased as the cells became more polarized. However, the recoil expansion rate of Y27632-treated cells was slightly larger than the recoil expansion rate of untreated cells at day 3 (Fig. 4H). The tortuosity of tight junctions in Y27632-treated MDCK cells was 1.02 (*N* = 29), suggesting a smaller medio-apical tension than in untreated MDCK cells based on our hypothesis. The larger recoil expansion rates of Y27632-treated cells can be explained by a decrease in cellular viscosity when cells are treated with Y27632 (19) since the recoil dynamics can be modelled as a viscoelastic system (23).

### Finding scutoid cells in a flat MDCK sheet is not predicted by the current vertex model

We noticed that cells within a cell monolayer changed neighbors along the apico-basal axis (Fig. 5A-C, highlighted cells; Movie S5), a characteristic of scutoids (4). Scutoids were originally predicted to arise from the curved geometry of cells experiencing strong circumferential tension (4). Gomez-Galvez *et al*. reported that scutoids do not exist in a flat cell sheet, in which cell-cell interactions consist of simple linear tension (4).

**Fig. 5.**
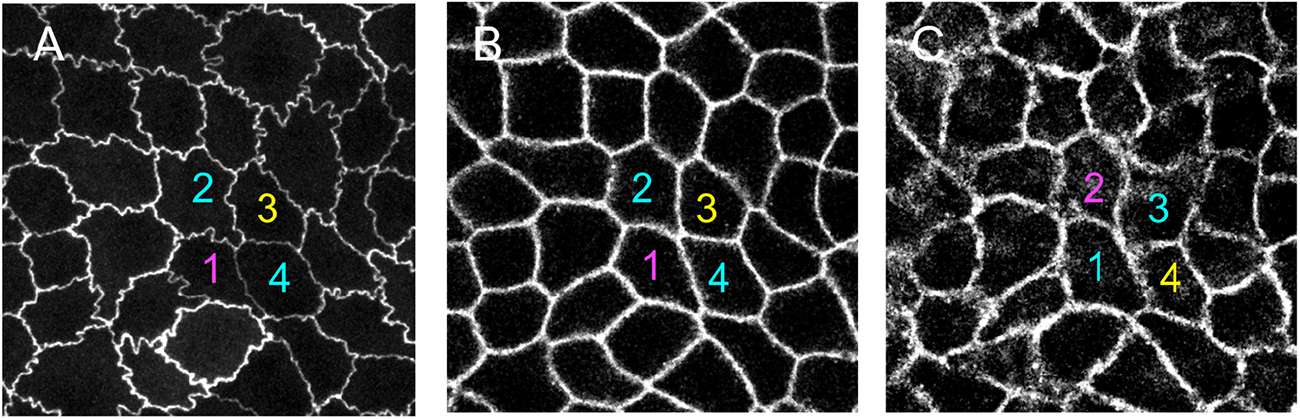
Scutoids exist on a flat substrate. (A) Apical cell shape is defined as the apical maximum projection of images of tight junctions. Here, cells 2 and 4 are neighbors (cyan) and separate cells 1 (magenta) and 3 (yellow). (B) Mid-cell shapes come from images of membranes at mid-cell along the apico-basal axis. Here, cells 2 and 4 are still neighbors. (C) Basal cell shapes come from membrane images slightly above the filter. Here, cells 2 and 4 do not share an edge and cells 1 and 3 are neighbors. All four cells are scutoids (Movie S5).

In contrast, we observed scutoids in a flat sheet. In MDCK cells grown for 3 days, we found scutoids constituted 46.6 ± 0.031% of cells (337 scutoids out of 653 cells, Materials and Methods), a surprisingly high prevalence (4). When MDCK cells seeded at the same cell density and at the same time were treated with 20 mM Y27632 for 24 h, this frequency decreased to 30.3 ± 0.046% (200 scutoids out of 663 cells). Critically, our observation of scutoids in a flat MDCK cell sheet indicates that modifications to the original model (4) are necessary. Although our observation of scutoids is not direct evidence for the existence of medio-apical tension, it suggests that cell-cell interactions may go beyond line tension to fill the discrepancy between the theory and the experimental observation. Medio-apical tension exerted at the apical cell surface provides a reasonable explanation for finding scutoids on a flat geometry. Treating cells with Y27632 reduces medio-apical tension, thereby decreasing the occurrence of scutoids.

## Conclusions

Here, we sought to determine the roles of apical actin cytoskeleton network in generating and transmitting tension at the apical surfaces and tight junctions of MDCK epithelial cells. Our findings demonstrate that there is non-negligible in-plane apical tension from the apical network coupled to the tight junction; this tension affects tight junction morphology as well as overall epithelial cell shape (including the scutoid morphology). The purse-string model based on the circumferential actomyosin belt (5, 6) is insufficient to describe cell expansion after laser ablation of a spot on the cell surface. The purse-string model also fails to capture the existence of scutoids within a flat cell sheet. Note that our experiments do not rule out a possible connection between the tight junction and the circumferential actomyosin belt, but our data emphasize the importance of apical in-plane tension, which has been overlooked to date.

Our work provides a new approach for thinking about the direction of force on the tight junction, and a reinterpretation of previous results. Although we determined here that in-plane tension is related to the sub-apical actin meshwork, further investigations are needed to reveal how in-plane apical tension is generated and transmitted through tight junctions. We believe that the actomyosin network is most likely the contractile terminal web (29-31) and that there are direct connections between the tight junction and the terminal web associated with microvilli. However, other mechanisms of force generation and transmission may be at play; for example, Li *et al*. (32) recently reported that actin polymerization pushes against apical junctions, maintaining E-cadherin adhesions.

Conventionally, tight junctions have not been associated with force transmission (19). However, our results suggest that tight junctions transmit apical tension. The adaptor protein ZO-1 is the molecule most likely to transmit apical in-plane tension (33). Depletion of ZO-1 from epithelial cells results in linear tight junctions as well as a smaller and more bulging apical surface (16, 17, 34). We hypothesize a direct connection between tight junction proteins and the terminal web actin network; super-resolution microscopy will be necessary for direct observations, although previously transmission electron microscopy was used to detect linkages between intestinal adsorptive cell tight junctions and microvillus actin (35). Recent work showed that the phase separation of ZO-1 drives formation of tight junction (36), which are mechanosensitive in gastrulating zebrafish embryos (37). In-plane tension may play an important role in ZO-1 phase separation in MDCK.

Theoretical modeling is an indispensable tool for deciphering morphogenesis (38). One of the most prevalent and powerful models of tissue morphogenesis is the “vertex model”, which often simplifies cell-cell contacts as straight lines and treats a flat two-dimensional cell sheet as a 2D system (3, 39, 40). Our results show that cell-cell interactions in a 2D cell sheet change along the apico-basal axis; in this situation, a 2D cell sheet should be still modeled as a 3D system. The tortuous cell-cell contacts caused by medio-apical tensions—which we have uncovered here—constitute a new factor for further vertex modeling. Integrating all of these factors into a model may explain the existence of scutoids in the flat geometry used here and further our understanding of organ morphogenesis.

## Materials and Methods

### Cell culture

MDCK cell lines were constructed by transfecting EGFP-ZO1 into MDCK cells already stably expressing mCherry-H2B (a gift from Dr. Chin-lin Guo’s lab) and selecting via serial dilution of G418 (Gibco, 10131). MDCK cells were maintained at 37 °C and 5% CO_2_ in air in MEM (Gibco, 11095) supplemented with 10% fetal bovine serum (Hyclone, SH30071.03), 10 mM sodium pyruvate (Gibco, 11360), 100 IU/mL penicillin, and 100 mg/mL streptomycin (Gibco, 15140122).

To culture a polarized monolayer cell sheet, we followed a previous protocol (41) by culturing cells on the outer side of a transwell (Corning, 3470). Cells were usually seeded as 40 μL of a suspension of 4.25 × 10^5^ cells/mL per 6.5-mm transwell. Cells were cultured for 3 days before laser ablation, with culture medium exchanged every day. Transwells were mounted on a custom-made glass-bottom chamber for confocal imaging. For treatment with ROCK inhibitor, cell sheets were grown for 2 days and treated with medium containing 20 μM Y27632 (StemCell, 72305) for 1 day.

### Immunofluorescence

Cells were washed with HBSS containing calcium and magnesium (ThermoFisher) three times. For membrane staining, we used MemBrite Fix 640/660 (Biotium) before fixing. We found that the prestaining solution reduced the tortuosity of the tight junctions (data not shown), probably by affecting cell-cell interactions when prestaining solution reacts with membrane proteins. We treated cells with prestaining solution at 1:2000 dilution (half the concentration suggested by the manufacturer) for 5 min and followed the manufacturer’s instructions for the rest of the experiment. Afterward, cells were fixed with 4% formaldehyde for 10 min and washed with phosphate-buffered saline (PBS) three times. Fixed cells were stained with Alexa 405 plus phalloidin (1:200, ThermoFisher) and washed with PBS three times. For imaging, the transwell was directly mounted onto a custom-made glass-bottom dish immersed in 1% DABCO (Sigma-Aldrich) dissolved in PBS.

### Laser ablation and imaging

For most laser-ablation experiments, cells were imaged through a 40x/1.2 NA water objective on an LSM 510-Meta-NLO (Zeiss) and ablated with a Chameleon Ultra femtosecond laser (Coherent Inc.) tuned to 750 nm with a maximum power output of 3.1 W. The ablating region was set as a box of 1×1 pixels, with a dwelling time of 1.6 μs or 3.2 μs. We tested laser powers from 20% to 60%. Due to the availability of the instrument, the recoil expansion measurements are carried out an LSM 980 (Zeiss) with the same objective and femtosecond laser as LSM 510. The ablation was done in the spot mode at 50% power. Time-lapse movies of MDCK cell sheets were acquired with a 40X/1.2 NA water lens on a Leica microscope (DMI 6000) equipped with a CSU22 spinning disk confocal scan head (Yokogawa) and an EMCCD (Oxford Instruments, iXon 885); cells were maintained at 37 °C and 5% CO_2_ with an LCI stage-top incubator. At each time point, we acquired a z-stack with 1 μm spacing through various channels; we used the maximum projected image for analysis. Membrane-and phalloidin-stained cells were imaged at finer z-spacing of 300 nm with a 63x/1.3 NA water lens on the same spinning disk confocal microscope or with a 63x/1.3 NA water lens on the LSM 880 (Zeiss) microscope.

### Image processing and data analysis

For live-cell imaging and immunostaining, data from tight junctions and nuclei channels were maximum projected as a single image before segmentation. Details of image segmentation appear in the SI Materials and Methods. Briefly, cells were segmented with the watershed function in MATLAB (R2018a, The MathWorks). Tight-junction contours were subpixel-refined from the dams of the watershed results (Fig. S6A-D, SI Materials and Methods). Vertices of individual cells were extracted with custom-written code by going through all the points along the cell boundary where neighbors changed (Fig. S6D, green dots). We defined the polygonal perimeter as the sum of all end-to-end lengths between adjacent vertices of a cell (Fig. S6D, cyan line). The tortuosity of a cell was defined as the cell’s contour length divided by its polygonal perimeter. The scutoids were identified by finding cells which have changes of neighbors along the apico-basal axis. The area expansion rate was computed as the maximum slope of the area vs. time data smoothed by a Gaussian kernel (42) (Fig. 1C). The onset time of the expansion was defined by the cross point of two straight lines fit from the baseline of the original area and the fitted expansion line (Fig. 1C). All statistical tests were carried out in OriginPro (2018, OriginLab). Correlation analyses were based on Pearson’s correlation coefficient and two-group comparisons were based on a two-sample Kolmogorov-Smirnov test.

## Acknowledgments

KH thanks Prof. Jason Swedlow for suggesting the laser-ablation experiments and Prof. Yu-Chiun Wang, Prof. Chin-ling Guo, Prof. Javier Buceta, and Prof. James Nelson for helpful discussions on the mechanics of epithelial cells. We thank former lab members Sung-chi Su and Jau-yi Wu for building the MDCK cell line stably expressing EGFP-ZO1 and mCherry-H2B. We thank Prof. I-ping Tu and Cheng-yu Hung for the kernel smoothing tip and Prof. Mong-na Lo for help with statistical analysis. Ablation experiments were carried out in the Imaging Core Facility at Institute of Molecular Biology at Academia Sinica with the assistance of Shu-ping Lee. Some confocal images were taken at the Imaging Core Facility at the Neuroscience Program of Academia Sinica with the assistance of Dr. Ya-jen Cheng. This work was supported by the Career Development Award of Academia Sinica (CDA-103-M07) and MOST 106-2112-M-001-029.

