## Supplementary Information for "Apical Actin-myosin Network Regulates the Tight Junction of Polarized Madin-Darby Canine Kidney Cells"

### Supporting Information

SI Materials and Methods

**F-actin Intensity Ratio of the Apical and Basal Planes.** F-actin intensity was measured by averaging all pixel values of the fluorescence channel for phalloidin per slice (Fig. S3). The intensity vs. z-slice curve was first smoothed and then two peaks were found, one near the apical plane and one near the basal plane. We define $I_{\text{apical}}$ as the maximum peak value near the apical plane and $I_{\text{basal}}$ as the maximum peak value near the basal plane. The F-actin apical/basal ratio is defined as $I_{\text{apical}} / I_{\text{basal}}$.

**Tight Junction Contours with Subpixel Fit.** Tight junctions (Fig. S4A) were first segmented as the dams from the watershed results (Fig. S4B), connecting pixel-resolved points around a cell (Fig. S4C, blue line). In order to compute tortuosity more accurately, we traced tight junctions with subpixel resolution; we followed and modified the method described by Guan, Wang, and Granick (2). Refinement procedures included moving contour points to the Gaussian-fit centers from the cross-section intensity profiles through the mid-points between the initial points using Gpufit (3). When the Gaussian fit was not good (for example its center point was beyond the cross-sectional length), the original point was kept. Intermediate contours were used to connect new center points and some unchanged original points; intermediate contours were further resampled and smoothed. Refinement procedures were repeated 40 times. The final refined contour trace (Fig. S6C, red line) was much closer to the tight-junction images than the original watershed contour (Fig. S6C, blue line).

It is crucial to compute the contour length based on the points after the subpixel fit, which yields more accurate length measurements than the pixel-resolved connected lines from the watershed procedure. We validated our computational method by generating synthesized images of circles as our ground-truth model (Fig. S6D). We generated circles of radii from 50 pixels to 200 pixels, consistent with typical MDCK image sizes, and calculated the percentage error of the mean pixel-resolved perimeters ($N = 10$) and subpixel-resolved perimeters ($N = 10$) from the ground truth (Fig. S6E). The larger the circle, the more accurate the subpixel-resolved perimeter (Fig. S6E). Based on our simulated results, the error of calculated perimeters is much less than 1%.


1. Schmidt U, Weigert M, Broaddus C, Myers G, editors. Cell Detection with Star-Convex Polygons2018; Cham: Springer International Publishing.
2. Guan J, Wang B, Granick S. Automated Single-Molecule Imaging To Track DNA Shape. Langmuir. 2011;27(10):6149-54.
3. Przybylski A, Thiel B, Keller-Findeisen J, Stock B, Bates M. Gpufit: An open-source toolkit for GPU-accelerated curve fitting. Sci Rep. 2017;7(1):15722.
4. Bi D, Lopez JH, Schwarz JM, Manning ML. A density-independent rigidity transition in biological tissues. Nat Phys. 2015;11(12):1074-9.
5. Fletcher Alexander G, Osterfield M, Baker Ruth E, Shvartsman Stanislav Y. Vertex Models of Epithelial Morphogenesis. Biophys J. 2014;106(11):2291-304.
6. Alt S, Ganguly P, Salbreux G. Vertex models: from cell mechanics to tissue morphogenesis. Philosophical Transactions of the Royal Society B: Biological Sciences. 2017;372(1720):20150520.
7. Wen F-L, Wang Y-C, Shibata T. Epithelial Folding Driven by Apical or Basal-Lateral Modulation: Geometric Features, Mechanical Inference, and Boundary Effects. Biophys J. 2017;112(12):2683-95.

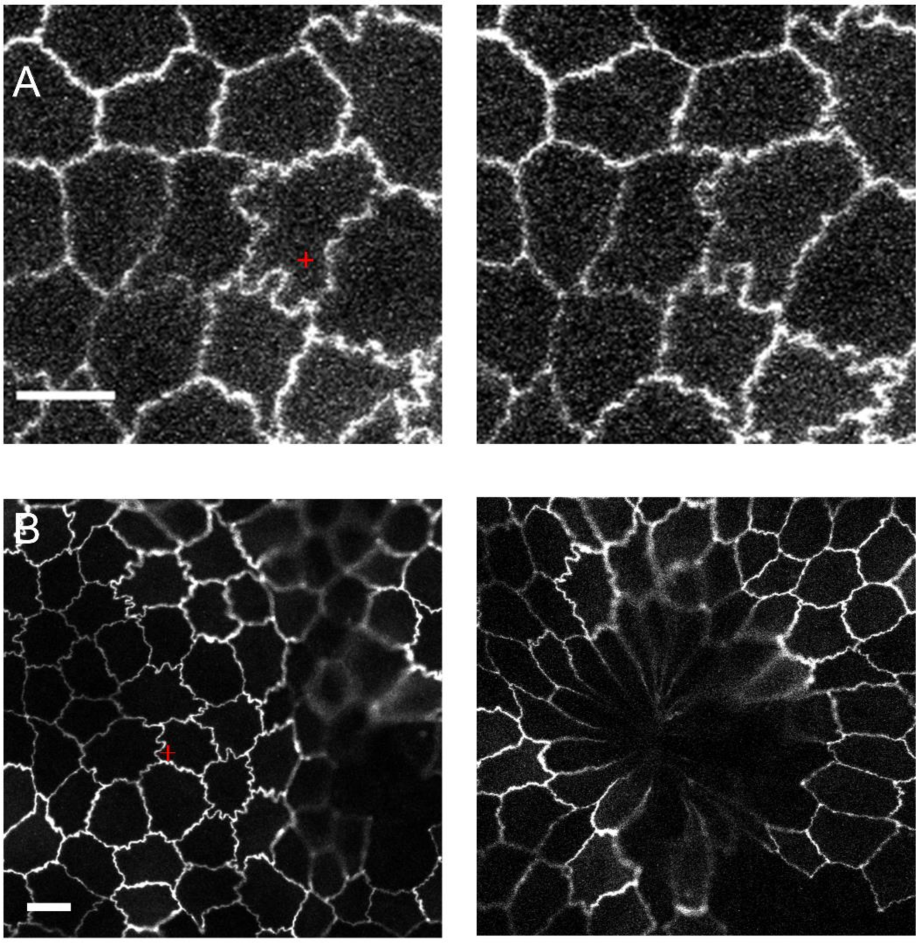


**Fig. S1.** Cells survive at low ablation power and die at high ablation power. (A) Confocal imaging of an MDCK cell sheet was carried out before ablation (left) and ~1 h after ablation (right) at (A) 32% laser power and (B) 60% power. The cross indicates the ablation site. Scale bar, 10 μm.

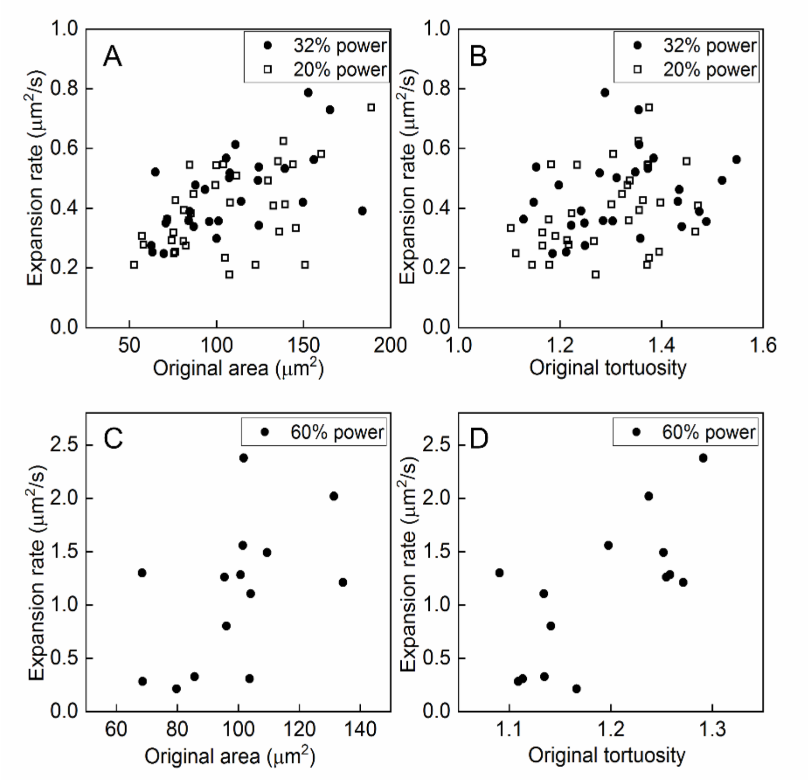


**Fig. S2.** Scatter plots of expansion rate vs. original apical surface area and tight-junction tortuosity. (A) After ablation, expansion rates and original areas are highly correlated for 32% (Pearson's $R$ = 0.81, $p$ = $10^{-10}$; $N$ =28) and 20% ($R$ = 0.51, $p$ = 0.002; $N$ =35) laser power. (B) Before ablation, expansion rates and original tortuosity are less correlated for 32% ($R$ = 0.22, $p$ = 0.24; $N$ = 28) and 20% ($R$ = 0.39, $p$ = 0.02; $N$ = 39) power. (C) Expansion rates and original areas are correlated after ablation at 60% power ($R$ = 0.51, $p$ = 0.06; $N$ = 14). (D) Expansion rates are correlated with tortuosity ($R$ = 0.6, $p$ = 0.09; $N$ = 14).

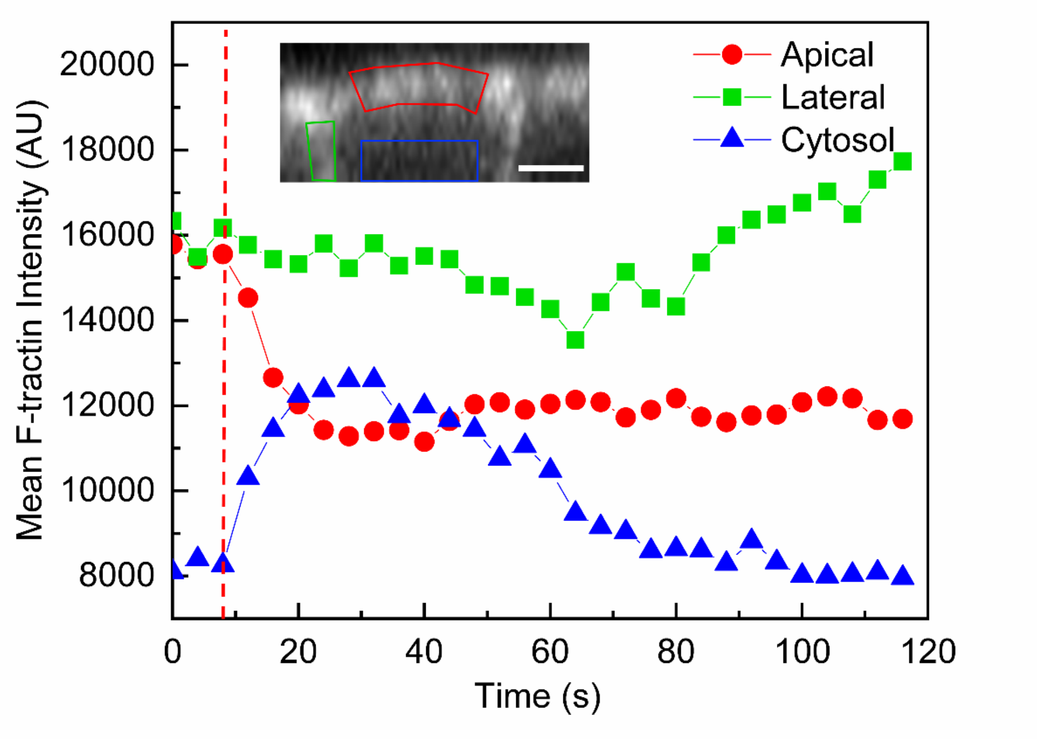


**Fig. S3**. Mean pixel intensity of F-tractin signals in the drawn regions of interest (ROIs) representing apical (red), lateral (green), and cytosol (blue) regions. The average apical F-tractin intensity decreases and the cytosol F-tractin intensity increases at similar rate after ablation (red dashed line). The apical F-tractin signal remains low and the cytosol F-tracin signals stay plateau for half a minute and decrease. The lateral F-tractin signals decrease slightly after ablation but increase when cytosol F-tractin signal decrease. (Inset) XZ slice of an MDCK cell expressing F-tractin with drawn ROIs. Scale bar, 3 μm.

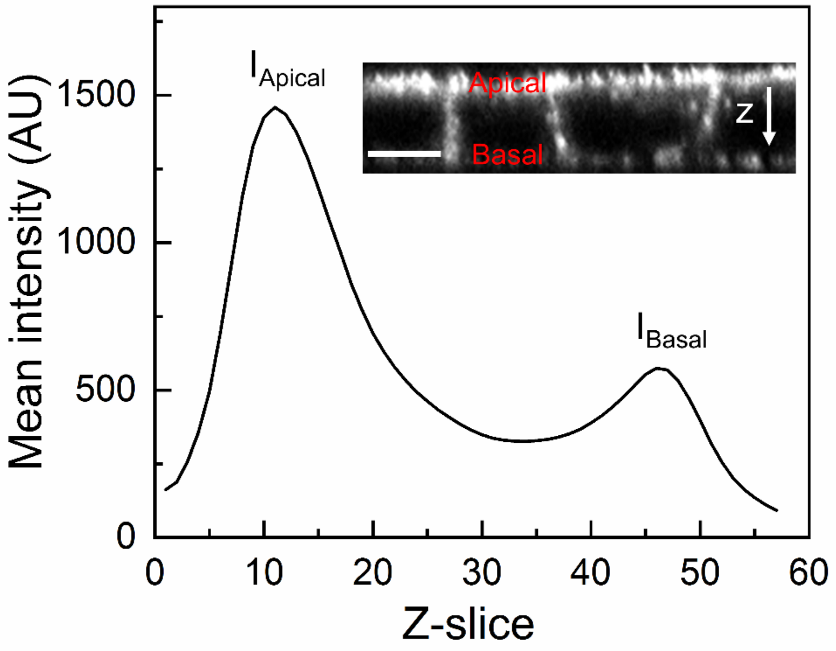


**Fig. S4**. Mean pixel intensity of phalloidin signal along the z-axis of MDCK cell sheet. (Inset) *xz* slice of an MDCK cell sheet stained with Alexa 405+ phalloidin. Scale bar, 5 µm.

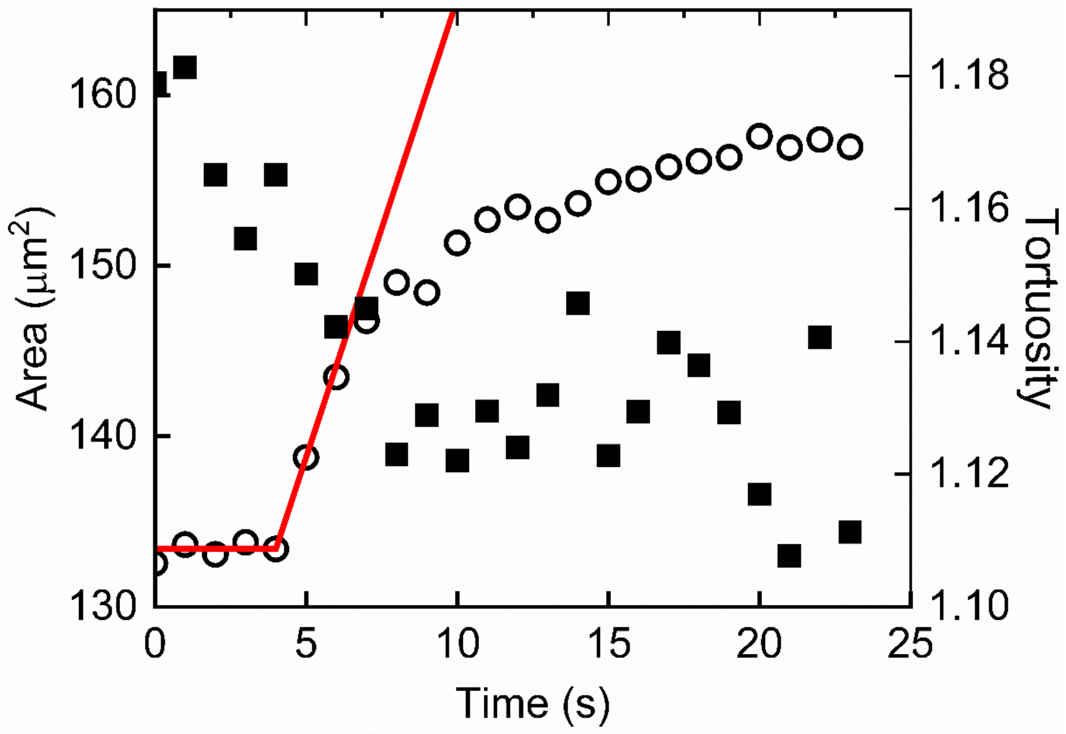


**Fig. S5.** Apical area increases sharply and tortuosity decreases when cells are ablated (red dashed line) at 50% laser power by the spot mode using LSM 980.

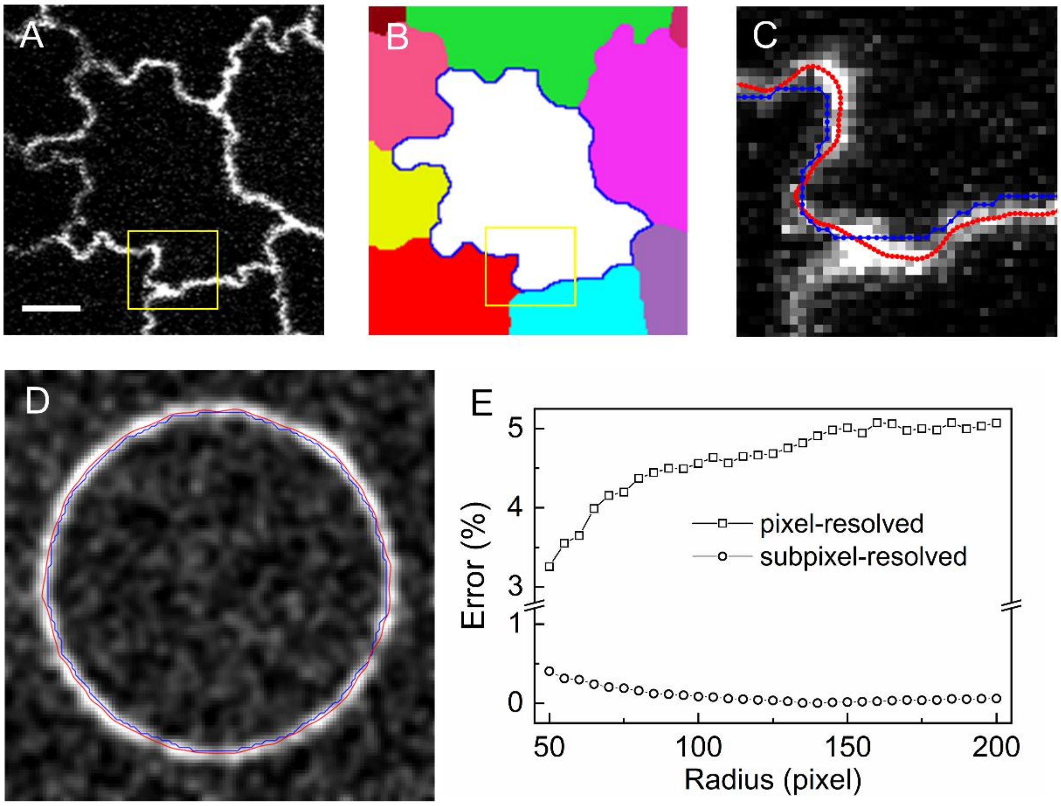


**Fig. S6.** Segmentation of cell morphology. (A) Raw image of tight junction. Scale bar, 5 μm. (B) Watershed image with the blue boundary as the initial pixel-resolved cell boundary, which is the dam from watershed. (C) Zoomed-in view (yellow box from (B)) of cell contours of a pixel-resolved dam (blue) and a subpixel-resolved curve (red) overlaid on an intensity image. (D) The ground truth model is a synthesized image of a circle, 50 pixels in radius. In this image, the pixel-resolved perimeter (blue) is 325.22 pixel, a 2.5% over-estimate, and the sub-pixel resolved perimeter is 314.515 pixel, constituting ~0.1% error. (F) Radius versus percentage error of pixel-resolved perimeters and subpixel-resolved perimeters deviating from ground-truth values. Each point is the average of 10 synthesized images with random noise and intensity fluctuations. The larger the radius, the larger the error from a pixel-resolved perimeter.

**Movie S1.** Ablating the apical surface of MDCK cells expressing GFP-ZO-1 at 32% laser power. The ablated point is marked with a cross. The apical surface of the cell expands gradually after ablation. Neighboring cells are also affected by ablation.

**Movie S2.** Ablating the mid-plane of MDCK cells expressing GFP-F-tractin at 32% laser power. The ablated point is marked with a cross. Scale bar, 3 μm. No changes in GFP-F-tractin are evident after ablation at the mid-section of the cell.

**Movie S3.** En face view of ablating the apical surface of MDCK cells expressing GFP-F-tractin at 32% laser power. The ablated point is marked with a cross. Scale bar, 5 μm.

**Movie S4**. Time-lapse movie of an MDCK cell sheet expressing GFP-ZO-1 from day 2 to day 3 at intervals of 10 minutes. Scale bar, 10 μm.

**Movie S5**. Apico-basal scan of an MDCK cell sheet stained with MemBrite Fix 640/660. Red curves outline cell boundaries and there is an intercalation of cell boundaries along the apico-basal axis. The movie further reconstructs cells with a rendering view. The exploding view of cells shows the internal vertex along the apico-basal axis; each cell is a scutoid.
